# SARS-CoV-2 RNA Persists in the Central Nervous System of Non-Human Primates Despite Clinical Recovery

**DOI:** 10.1101/2023.08.29.555368

**Authors:** Hailong Li, Kristen A. McLaurin, Charles F. Mactutus, Jay Rappaport, Prasun K. Datta, Rosemarie M. Booze

## Abstract

Adverse neurological and psychiatric outcomes, collectively termed the post-acute sequelae of SARS-CoV-2 infection (PASC), persist in adults clinically recovered from COVID-19. Effective therapeutic interventions are fundamental to reducing the burden of PASC, necessitating an investigation of the pathophysiology underlying the debilitating neurological symptoms associated with the condition. Herein, eight non-human primates (Wild-Caught African Green Monkeys, *n*=4; Indian Rhesus Macaques, *n*=4) were inoculated with the SARS-CoV-2 isolate USA-WA1/2020 by either small particle aerosol or via multiple routes. At necropsy, tissue from the olfactory epithelium and pyriform cortex/amygdala of SARS-CoV-2 infected non-human primates were collected for ribonucleic acid *in situ* hybridization (i.e., RNAscope). First, angiotensin-converting enzyme 2 (ACE2) and transmembrane serine protease 2 (TMPRSS2) mRNA are downregulated in the pyriform cortex/amygdala of non-human primates clinically recovered from SARS-CoV-2 inoculation relative to wildtype controls. Second, abundant SARS-CoV-2 mRNA was detected in clinically recovered non-human primates; mRNA which is predominantly harbored in pericytes. Collectively, examination of post-mortem pyriform cortex/amygdala brain tissue of non-human primates clinically recovered from SARS-CoV-2 infection revealed two early pathophysiological mechanisms potentially underlying PASC. Indeed, therapeutic interventions targeting the downregulation of ACE2, decreased expression of TMPRSS2, and/or persistent infection of pericytes in the central nervous system may effectively mitigate the debilitating symptoms of PASC.

## Dear Editor

The post-acute sequelae of severe acute respiratory syndrome coronavirus 2 (SARS-CoV-2) infection is a multisystemic condition characterized by the persistence or emergence of COVID-19 symptoms months after acute infection. Neurological symptoms, including neurocognitive impairments, neuropsychiatric disturbances, and olfactory dysfunction, are reported by a significant proportion (e.g., 31.3%, 21.1%, and 15.3%, respectively^1-2^) of individuals months after acute SARS-CoV-2 infection; symptoms which adversely impact working capacity^1^. Effective therapeutic interventions are fundamental to reducing the burden of post-acute sequelae of SARS-CoV-2 (PASC), necessitating an investigation of the pathophysiology underlying the debilitating neurological symptoms associated with the condition.

Preclinical biological systems are fundamental to the comprehensive investigation of the pathogenesis of PASC following clinical recovery. Non-human primates, in particular, are naturally susceptible to SARS-CoV-2, whereby species-specific differences in SARS-CoV-2 susceptibilities have been reported^3^. Therefore, in the present study, eight non-human primates (Aged (16 Years Old), Wild-Caught African Green Monkeys, *n*=4; Indian Rhesus Macaques (13-15 Years Old), *n*=4) were inoculated with the SARS-CoV-2 isolate USA-WA1/2020 by either small particle aerosol (Dose of 1 x 10^4^ Plaque-Forming Units) or via multiple routes (i.e., oral, nasal, intratracheal and conjunctival; Cumulative Dose of 3.61 x 10^6^ Plaque-Forming Units^4^). Comprehensive details on the clinical phenotype observed in these animals are reported by Blair et al.^4^. In brief, high levels of SARS-CoV-2 RNA, extracted from mucosal swab and bronchial brush samples, were detected in both African green monkeys (AGMs) and Rhesus macaques (RMs); albeit no statistically significant species or dose effects were observed. Two AGMs (AGM1, AGM2) developed acute respiratory distress syndrome, necessitating humane euthanasia eight and twenty-two days post-infection, respectively. The six remaining primates (AGM: *n*=2; RM: *n*=4) clinically recovered from SARS-CoV-2 inoculation, whereby viral loads were undetectable, and no significant clinical pathology was observed at the termination of experimentation (i.e., 21-28 Days Post Infection). Taken together, both AGMs and RMs recapitulate key aspects of the clinical syndrome, including clinical recovery (in six of the primates) and heterogeneity in disease phenotype (i.e., Mild to Severe Disease), affording a biological system to investigate early pathophysiological changes underlying the neurological symptoms associated with PASC.

At necropsy, tissue from the olfactory epithelium and pyriform cortex/amygdala of SARS-CoV-2 infected primates were collected for further examination. The olfactory epithelium, which is involved in the transduction of olfactory information, lines the olfactory cleft of the nasal cavity. This neuroanatomical location may render it uniquely vulnerable to invasion by SARS-CoV-2 infection. From a functional perspective, the olfactory epithelium transmits odorant information to the pyriform cortex, where it is processed. The pyriform cortex is reciprocally connected to brain regions involved in emotional processing (e.g., amygdala) and cognitive function (e.g., orbitofrontal cortex). Hence, the characteristics (i.e., neuroanatomical location, function, neural connectivity) of the olfactory epithelium and pyriform cortex/amygdala necessitate considering these brain regions in the early pathophysiological changes underlying PASC.

An innovative ribonucleic acid (RNA) *in situ* hybridization approach (RNAscope^5^), which utilizes a unique probe design to concurrently amplify target-specific signals and suppress background noise, was first employed to investigate the expression of SARS-CoV-2 entry proteins (i.e., Angiotensin-Converting Enzyme 2 (ACE2), Transmembrane Serine Protease 2 (TMPRSS2)) in the pyriform cortex/amygdala of wildtype (Indian Rhesus Macaques; *n*=4) and clinically recovered SARS-CoV-2 non-human primates. A fluorescently labeled probe was utilized for direct visualization, whereby a punctate, single-dot staining pattern is routinely observed; each single-dot represents a single messenger RNA (mRNA) transcript^5^. Consistent with previous reports, both ACE2 (Figure 1A-B) and TMPRSS2 (Figure 1D-E) were highly expressed in the pyriform cortex/amygdala of both wildtype and SARS-CoV-2 inoculated non-human primates. A history of SARS-CoV-2 inoculation, however, induced a significant downregulation of both ACE2 [Figure 1C; *F*(1,11)=10.0, *p*≤0.01] and TMPRSS2 [Figure 1F; *F*(1,11)=7.5, *p*≤0.02] mRNA in the pyriform cortex/amygdala. The downregulation of ACE2, in particular, likely results in the exacerbation of inflammatory responses via the overactivation of Angiotensin II (for review^6^); the physiological implications of TMPRSS2 downregulation in the central nervous system, however, require further study.

**Figure 1.**
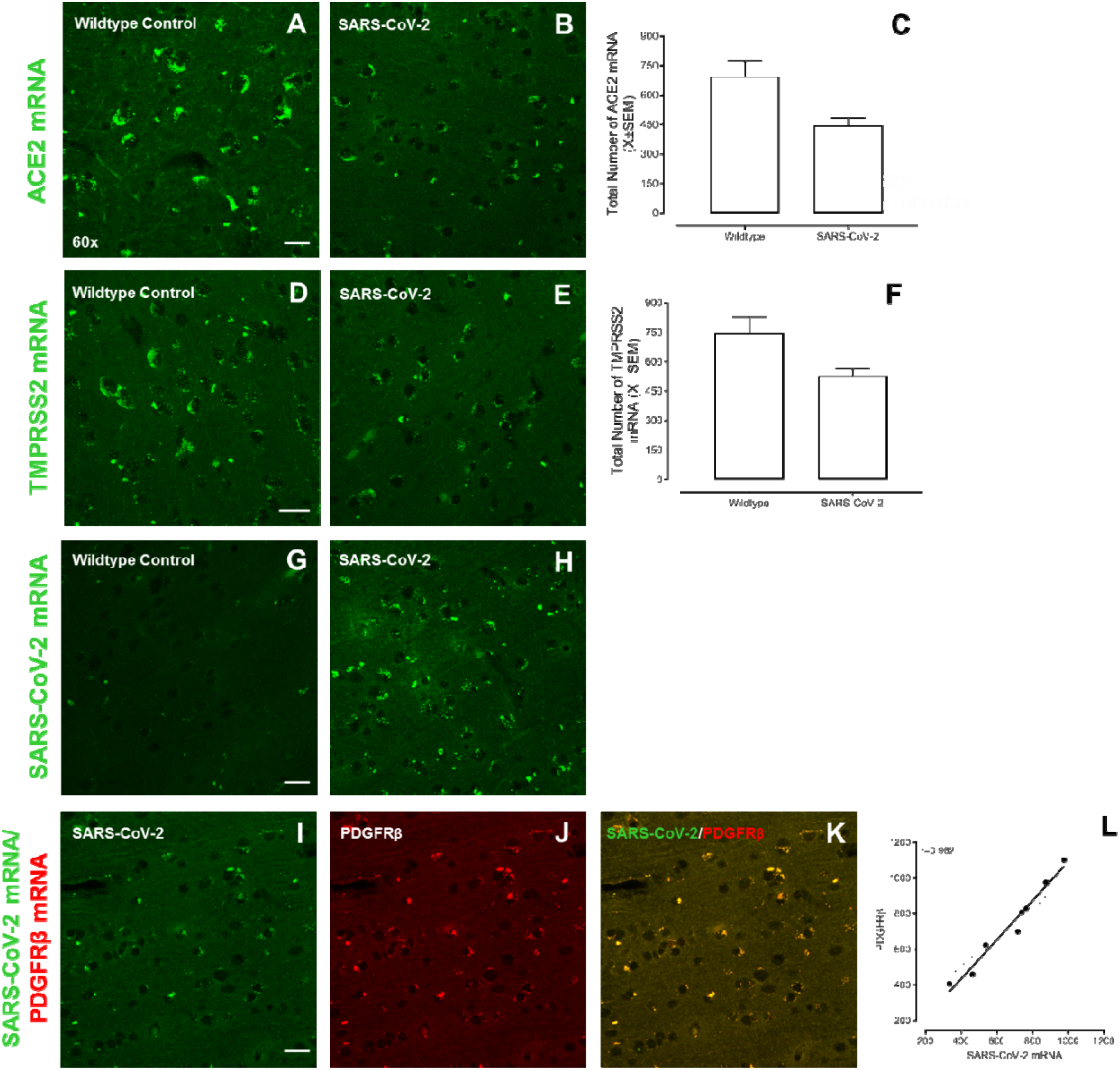
Eight non-human primates (Aged (16 Years Old), Wild-Caught African Green Monkeys, *n*=4; Indian Rhesus Macaques (13-15 Years Old), *n*=4) were inoculated with the SARS-CoV-2 isolate USA-WA1/2020 by either small particle aerosol or via multiple routes (i.e., oral, nasal, intratracheal and conjunctival^4^). After establishing clinical recovery (i.e., undetectable SARS-CoV-2 viral load, no significant clinical pathology), brain tissue from the pyriform cortex/amygdala was collected for analysis. Brain tissue from historic wild-type non-human primates (i.e., Rhesus Macaques; *n*=4) was also examined. Representative confocal images are shown from the pyriform cortex/amygdala of wildtype and clinically recovered non-human primates, as appropriate. Angiotensin-converting enzyme 2 (ACE2; **A-B**) and Transmembrane Serine Protease 2 (TMPRSS2; **D-E**), two integral SARS-CoV-2 entry proteins, were observed in the pyriform cortex/amygdala affording a venue by which SARS-CoV-2 may enter the central nervous system. Furthermore, a history of SARS-CoV-2 infection resulted in a statistically significant downregulation of both ACE2 (**C**) and TMPRSS2 (**F**). SARS-CoV-2 mRNA does indeed persist in the pyriform cortex (**H**) of non-human primates despite clinical recovery; no prominent fluorescent SARS-CoV-2 mRNA were observed in the pyriform cortex/amygdala of non-infected primates (**G**). The predominant cell type expressing SARS-CoV-2 was subsequently investigated in clinically recovered non-human primates, whereby high co-localization between SARS-CoV-2 mRNA and platelet-derived growth factor receptor beta (PDGFRβ), a marker for pericytes, was observed (**I-L**). Scale bar, 10 µm.

In addition, expression of ACE2 and TMPRSS2 in the brain affords a venue through which SARS-CoV-2 may enter the central nervous system. Indeed, SARS-CoV-2 RNA has been observed in autopsy brain tissues of patients who died with COVID-19 ^e.g., 7^; the expression of SARS-CoV-2 in the brain following clinical recovery (i.e., undetectable SARS-CoV-2 viral loads), however, has not been systematically evaluated. To address this knowledge gap, RNAscope was performed in conjunction with a highly specific probe for SARS-CoV-2 mRNA to evaluate the persistence of viral mRNA in the central nervous system. Abundant SARS-CoV-2 mRNA was detected in the pyriform cortex/amygdala (Figure 1H) of clinically recovered non-human primates. As expected, no prominent fluorescent SARS-CoV-2 mRNA was observed in the pyriform cortex/amygdala of wild-type non-primates (Figure 1G). Notably, ACE2, TMPRSS2, and SARS-CoV-2 mRNA were also observed in the olfactory epithelium, albeit at lower levels than the pyriform cortex/amygdala (Supplementary Figure 1); observations which support the persistence of SARS-CoV-2 in the central nervous system more broadly.

The persistence of SARS-CoV-2 mRNA in the brains of clinically recovered non-human primates necessitates an investigation of the cell type being infiltrated by the virus. Brain pericytes, in particular, abundantly co-express the SARS-CoV-2 cellular entry receptors ACE2 and TMPRSS2^8,^ affording a venue through which SARS-CoV-2 may enter these cells. To evaluate the hypothesis, brain tissue from the pyriform cortex/amygdala was dual-labeled for SARS-CoV-2 mRNA and platelet-derived growth factor receptor beta (PDGFRβ), a biomarker exclusively expressed in pericytes in the adult brain^9^. SARS-CoV-2 mRNA and PDGFRβ exhibited high co-localization (*r* = 0.982) in all experimental primates (Figure 1I-L). Pericytes, therefore, harbor SARS-CoV-2 in the central nervous system of non-human primates despite full clinical recovery; observations which extend upon previous reports documenting the productive infection of pericyte-like cells within cortical organoids^10^. Thus, SARS-CoV-2 infection of pericytes may lead to the central nervous system (CNS) manifestations of COVID-19, including inflammation and pericytes-mediated blood flow reductions.

Given the persistence of SARS-CoV-2 mRNA in the brain, advanced statistical approaches were utilized to evaluate whether the acute clinical disease phenotype was predictive of chronic brain infection in the pyriform cortex/amygdala. Specifically, regression analyses were conducted to evaluate whether viral loads and/or clinical symptomology were associated with the total number of SARS-CoV-2 mRNA (Supplementary Figure 2). There was no statistically significant relationship (H_0_: β_1_=0; *p*>0.05) between viral loads (from buccal, nasal, pharyngeal, bronchial brush, etc.), clinical assessment, or lung histopathologic score and the total number of SARS-CoV-2 mRNA in the pyriform cortex/amygdala. Thus, the acute clinical disease phenotype doesn’t predict the extent to which SARS-CoV-2 mRNA invades the central nervous system.

Taken together, examination of post-mortem pyriform cortex/amygdala brain tissue of non-human primates clinically recovered from SARS-CoV-2 infection revealed two early pathophysiological mechanisms potentially underlying PASC. First, a history of SARS-CoV-2 inoculation results in the downregulation of ACE2 and TMPRSS2 mRNA in the pyriform cortex/amygdala. Second, SARS-CoV-2 mRNA are harbored in pericytes of non-human primates clinically recovered from SARS-CoV-2 infection. Additional studies are necessary to identify long-term CNS alterations induced by SARS-CoV-2, as brain tissue was collected from non-human primates only 28 days after inoculation. Therapeutic interventions targeting the downregulation of ACE2, decreased expression of TMPRSS2, and/or persistent infection of pericytes in the central nervous system may effectively mitigate the debilitating symptoms of PASC.

## DATA AVAILABILITY

Supplemental information and raw data are available from the corresponding authors.

## ACKNOWLEDGMENTS

This work was supported in part by grants from NIH (National Institute on Drug Abuse, R01-DA013137; National Institute on Drug Abuse, K99-DA056288; National Institute on Drug Abuse, R01-059310; National Institute of Mental Health, R01-MH106392; National Institute of Neurological Disorders and Stroke, R01-NS100624; NIH Office of the Director, P51OD011104-59). Dr. Kristen A. McLaurin is now an Assistant Professor in the College of Pharmacy (Department of Pharmaceutical Sciences) at the University of Kentucky.

## AUTHOR CONTRIBUTIONS

R.M.B and C.F.M conceptualized the study. H.L. performed the experiments. K.A.M and H.L. analyzed the data. H.L., K.A.M., P.K.D., R.M.B., and C.F.M. wrote the manuscript. J.R. and P.K.D. provided the non-human primate samples. All authors have read and agreed to the published version of the manuscript.

