## Supplementary Information for "SARS-CoV-2 RNA Persists in the Central Nervous System of Non-Human Primates Despite Clinical Recovery"

[1]Cognitive and Neural Science Program
Department of Psychology
Barnwell College
University of South Carolina
1512 Pendleton Street
Columbia, SC 29208

[2]Tulane National Primate Research Center
18703 Three Rivers Road
Covington, LA 70433

**Address correspondence to:**

Rosemarie M. Booze, Ph.D.
Carolina Trustees Professor and Bicentennial Endowed Chair of Behavioral Neuroscience
Department of Psychology
1512 Pendleton Street
University of South Carolina
Columbia, SC 29208


**This PDF file includes:**

Materials and Methods

Figs. S1 to S2

Tables S1 to S3

### Materials and Methods

Study Approval

The experimental protocol #P0435 was approved by The Institutional Animal Care and Use Committee of Tulane University. The Tulane National Primate Research Center is fully accredited by the American Association for Accreditation of Laboratory Animal Care. Handling, inactivation, and removal of the non-human primate tissue from BSL3 containment was approved by The Tulane University Institutional Biosafety Committee. All experimental procedures handled in University of South Carolina, were approved by institutional biosafety committee of University of South Carolina (#300268).

Nonhuman primate SARS-CoV-2 infection model

The African Green Monkeys (AGMs) were wild caught in St. Kitts and Nevis and supplied by a USA importer under permit number 524/2019 dated 22/01/2019 (CITES Management Authority, Ministry of Agriculture, St. Kitts and Nevis) and U.S Fish and Wildlife Services (Import License number 110542 dated 01/29/2019, and housed at Tulane National Primate Research Center, Covington, for this study. Eight non-human primates (Aged (16 Years Old), AGMs $n$=4; Indian Rhesus Macaques (13-15 Years Old), $n$=4) were inoculated with the SARS-CoV-2 isolate USA-WA1/2020 (MN985325.1). Details regarding the preparation and confirmation of the virus stock are available in Blair et al[1].

RNAscope *in situ* Hybridization

RNAscope *in situ* hybridization was used to detect the expression of SARS-CoV-2, ACE2, and TMPRSS2 mRNA in the olfactory epithelium and pyriform cortex/amygdala of wildtype and clinically recovered non-human primates. The RNAscope *in situ* hybridization protocol utilized in the present study was described in detail by Li et al[2], albeit with minor modifications. Briefly, the olfactory epithelium and pyriform cortex/amygdala were fixed and embedded in paraffin. 5 µm sections were cut using a microtome and mounted onto SuperFrost Plus slides. After the sections were dried at 60°C for one hour, they were submerged in xylene (10 minutes, ×2) and 100% ethanol (10 minutes, ×2). Sections were subsequently boiled in Target Retrieval reagent for 15 minutes. RNA *in situ* hybridization was subsequently conducted using the RNAscope Multiplex Fluorescent Assay (Advanced Cell Diagnostics, Inc., Newark, CA, USA), whereby sections were hybridized with a specific probe for ACE2, TMPRSS2, or SARS-CoV-2 mRNA (see Supplementary Table S1). Following all amplification steps, slides were mounted with Pro-Long Gold Antifade (Invitrogen, Carlsbad, CA), cover-slipped, and stored at 4°C in the dark. Z-stack images were obtained using a 60× objective on a Nikon TE-2000E confocal microscope utilizing Nikon's EZ-C1 software (version 3.81b).

Quantification and Statistical Analyses

For the quantification of the total number of SARS-CoV-2 mRNA in the pyriform cortex/amygdala, the number of dots was quantified by two independent experimenters (Interrater Reliability: $r$=0.847). The total number of SARS-CoV-2 mRNA in the olfactory epithelium (i.e., sustentacular layer, basal stem cell layer) and the evaluation of co-localization were conducted by one experimenter.

GraphPad Prism 5 (GraphPad Software Inc., La Jolla, CA, USA) was used to create figures and conduct regression analyses. A $p \leq 0.05$ was considered statistically significant for all analyses. Analysis of variance techniques (SPSS Statistics 29, IBM Corporation, Somer, NY, USA) were utilized to evaluate how a history of SARS-CoV-2 inoculation impacts ACE2 and TMPRSS2 mRNA. Exposure (Wildtype Control vs. SARS-CoV-2 Inoculation) served as the between-subjects factor.

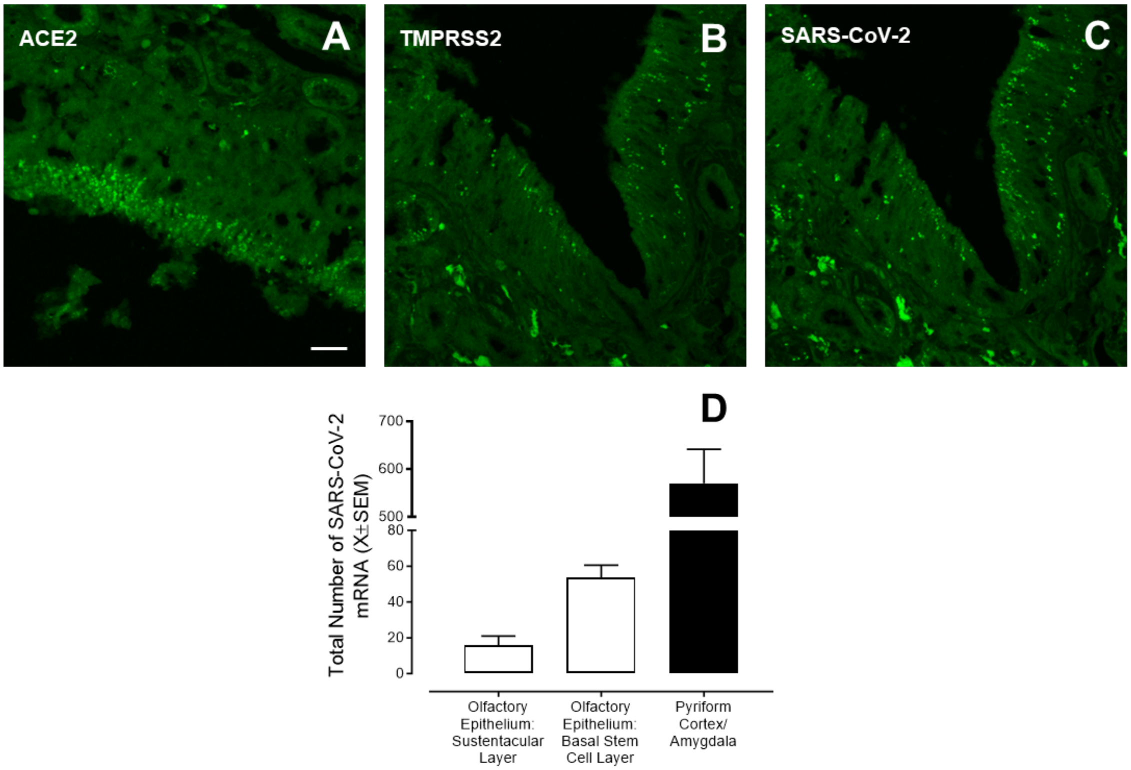


**Figure S1. Angiotensin-converting enzyme 2 (ACE2; A), Transmembrane Serine Protease 2 (TMPRSS2; B), and SARS-CoV-2 mRNA (C) were also observed in the olfactory epithelium of clinically recovered SARS-CoV-2 infected non-human primates.** Representative confocal images are shown from the olfactory epithelium of clinically recovered non-human primates. The total number of SARS-CoV-2 mRNA in the olfactory epithelium (Sustentacular Layer and Basal Stem Cell Layer) and pyriform cortex/amygdala were quantified by counting the number of fluorescent green dots, whereby each dot represents a single mRNA transcript[5] (**D**). The pyriform cortex/amygdala harbored a significantly greater number of SARS-CoV-2 mRNA than the olfactory epithelium.

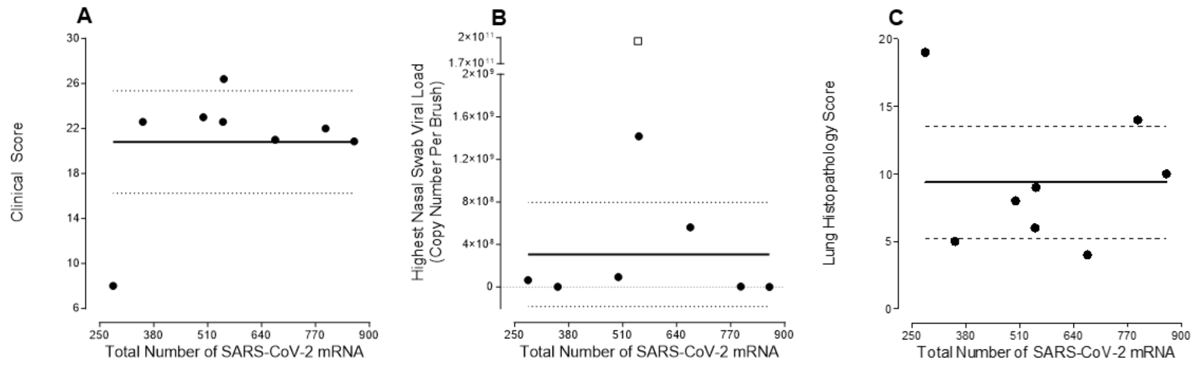


**Figure S2.** The acute clinical disease phenotype doesn't predict the extent to which SARS-CoV-2 mRNA invades the central nervous system. Regression analyses revealed no statistically significant relationship ($H_0$: $\beta_1=0$; $p>0.05$) between the clinical assessment (**A**), highest measured nasal swab viral load (**B**), or lung histopathology score (**C**) and the total number of SARS-CoV-2 mRNA in the pyriform cortex/amygdala. The square marker in (**B**) was an outlier, defined as greater than two standard deviations away from the mean, and was not included in the regression analyses.

**Table S1. Probes for RNAscope *in situ* assay**

| Probe | Cat. No. | Color | Comments |
|---|---|---|---|
| RNAscope Probe - V-nCoV2019-S | 848561, ACDbio | Green | SARS-CoV-2 mRNA |
| RNAscope Probe - Hs-ACE2 | 848151, ACDbio | Green | ACE2 |
| RNAscope Probe - Hs-TMPRSS2 | 470341, ACDbio | Green | TMPRSS2 |
| RNAscope Probe - Mfa-PDGFRB-C2 | 545251-C2, ACDbio | Red | PDGFRβ (Pericytes) |
| RNAscope 3-plex Negative Control | 320871 | | DapB (*Bacillus subtilis* strain) |
| RNAscope 3-plex Positive Control | 320901 | | Mfa-Polr2a |

**Table S3. Animal information including species, source, route of exposure, and demographic information from each animal in the study. Additional information has been reported by Blair et al.[4].**

| ID | Species | Source | Age (Yr) | Sex | Weight (kg) | Exposure (Dose) |
|---|---|---|---|---|---|---|
| AGM1 | *Chlorocebus aethiops sabaeus* | Wild caught, St. Kitts | 16 | F | 4.3 | Aerosol ($1x10^4$ PFU) |
| AGM2 | *Chlorocebus aethiops sabaeus* | Wild caught, St. Kitts | 16 | F | 3.9 | Multiroute ($3.61x10^6$ PFU) |
| AGM3 | *Chlorocebus aethiops sabaeus* | Wild caught, St. Kitts | 16 | M | 6.9 | Multiroute ($3.61x10^6$ PFU) |
| AGM4 | *Chlorocebus aethiops sabaeus* | Wild caught, St. Kitts | 16 | M | 7.5 | Aerosol ($1x10^4$ PFU) |
| RM1 | *Macaca mulatta* | Born at TNPRC | 14 | M | 16.7 | Multiroute ($3.61x10^6$ PFU) |
| RM2 | *Macaca mulatta* | Born at TNPRC | 13 | F | 6.9 | Multiroute ($3.61x10^6$ PFU) |
| RM3 | *Macaca mulatta* | Born at TNPRC | 13 | M | 11.6 | Aerosol ($1x10^4$ PFU) |
| RM4 | *Macaca mulatta* | Born at TNPRC | 15 | M | 11 | Aerosol ($1x10^4$ PFU) |
